# Designer DNA architecture offers precise and multivalent spatial pattern-recognition for viral sensing and inhibition

**DOI:** 10.1101/608380

**Authors:** Paul S. Kwon, Shaokang Ren, Seok-Joon Kwon, Megan E. Kizer, Lili Kuo, Feng Zhou, Fuming Zhang, Domyoung Kim, Keith Fraser, Laura D. Kramer, Nadrian C. Seeman, Jonathan S. Dordick, Robert J. Linhardt, Jie Chao, Xing Wang

**Affiliations:** Department of Chemistry & Chemical Biology, Center for Biotechnology and Interdisciplinary Studies, Rensselaer Polytechnic Institute, Troy, NY 12180, USA; Department of Neuroscience, Johns Hopkins University, Baltimore, MD, 21218, USA; Key Laboratory for Organic Electronics and Information Displays and Jiangsu Key Laboratory for Biosensors, Institute of Advanced Materials, National Synergetic Innovation Center for Advanced Materials, Nanjing University of Posts and Telecommunications, Nanjing 210023, China; Department of Chemical and Biological Engineering, Center for Biotechnology and Interdisciplinary Studies, Rensselaer Polytechnic Institute, Troy, NY 12180, USA; Wadsworth Center, New York State Department of Health, Albany, NY 12159, USA; Department of Chemistry, New York University, New York, NY 10003, USA; Center for Soft Matter Research, New York University, New York, NY 10003, USA

## Abstract

DNA, when folded into nanostructures of customizable shapes, is capable of spacing and arranging external ligands in a desired geometric pattern with nanometer-precision. This allows DNA to serve as an excellent, biocompatible scaffold for complex spatial pattern-recognizing displays. In this report, we demonstrate that a templated designer DNA nanostructure achieves multi-ligand display with precise spatial pattern-recognition, representing a unique strategy in synthesizing potent viral sensors and inhibitors. Specifically, a star-shaped DNA architecture, carrying five molecular beacon-like motifs, was constructed to display ten dengue virus envelope protein domain-III (ED3)-binding aptamers into a 2D pattern precisely matching the pentagonal arrangement of ED3 clusters on the dengue viral surface. The resulting spatial pattern recognition and multivalent interactions achieve high dengue-binding avidity, conferring direct, highly-sensitive, facile, low-cost, and rapid sensing as well as potent viral inhibition capability. Our molecular-platform design strategy could be adapted to detect and combat other disease-causing pathogens, including bacteria and microbial-toxins, by generating the requisite ligand patterns on customized DNA nanoarchitectures.

---

In viral detection, the gold standard diagnostic methods employ virus isolation, antigen or antibody capture immunoassays, and molecular diagnostic tests.^1–4^ These are typically time-consuming, expensive, and require a sophisticated clinical laboratory setup and technical expertise. Avoiding standard ELISA and RT-PCR based methods while achieving higher detection sensitivity with an easy, quick, and low-cost process would enable facile sensing for timely detection and control of pandemic outbreaks within surveillance and diagnostic networks; this would be particularly valuable in low-resource or field-applicable environments. In viral therapeutics, prevention or treatment of viral infection typically relies on neutralizing antibodies (NAbs) against target epitopes on the viral surface. Production of NAbs can be triggered by vaccines or active viruses that invade hosts. However, NAbs may induce antibody-dependent enhancement of infection (e.g., dengue^5^) or may not provide protection against new epidemics as a result of genetic drift (e.g., antigenic drift in influenza^6^). A customizable molecular scaffold can be generated to fit any surface pattern to recognize the drifting virus and serve as a more reliable platform for viral therapeutics. This may be particularly useful when fighting many World Health Organization priority “blueprint-viruses” such as Zika and HIV for which no approved or effective vaccines are available.

Infectious diseases, including viruses, bacteria, and microbial toxins, present a unique spatial pattern of antigens on their surfaces.^7–9^ Existing weakly binding ligands (called “binders” herein) that interact with these epitopes can be linked to a scaffold, but exhibit limited improvement on multivalent binding avidity when sporadically matching the average spacing of epitopes.^10–14^ Herein, we demonstrate a unique strategy for potent viral detection and inhibition through a precise, spatial pattern-recognizing display of weak binders in two-dimensional (2D) space. This approach is particularly important for targeting disease-causing pathogens with complex patterns of surface antigens such as dengue virus (DENV). Previously used synthetic scaffolds (polymers, dendrimers, nanofibers, nanoparticles and lipid nanoemulsions) have been toxic and cannot address these patterns, because they are not as precise in ligand spacing or provide limited control over the scaffold shape and ligand valency^15–18^ needed for the ideal and precise spatial display of 2D ligand patterns. Designer DNA nanostructures^19–23^ are able to overcome all these limitations because they can be designed into various nontoxic/biocompatible^24–27^ and biologically stable^27–31^ 2D or three-dimensional (3D) platforms capable of controlling external ligand valency and spatial arrangements. Most importantly, they are capable of displaying molecular ligands in complex 2D/3D geometric patterns with precise nanometer (nm) inter-ligand spacings.^20,32,33^ Such DNA-based nanostructures are biocompatible and exhibit some circulatory time up to 24 h,^27^ but will eventually be removed from the blood by the liver^27^ and kidney^34,35^ and thus, could serve as therapeutic agents to inhibit and clear the virus from the patient.

In this study, we demonstrate our unique molecular platform strategy by directly targeting DENV virions through a star-shaped DNA nanoarchitecture. This architecture was strategically chosen and designed to multivalently arrange molecularly recognizable binders in a 2D manner to precisely mimic the complex spatial pattern of the DENV viral envelope proteins. Nanometer-level precision matching of binders to protein epitope enables a potent interaction for sensing and inhibition.

Our viral sensing approach does not rely on nucleic acid amplification, sophisticated laboratory equipment, or other lengthy steps needed for viral isolation, ELISA, or RT-PCR based gold standard methods, yet provides much higher sensitivity (detection limit of 100 pfu/mL or 1,000 pfu/mL in human blood serum or plasma) through a much easier (simple mix-and-read), faster (within 5 minutes), and cost-effective (∼ $0.15 per test) assay. All these aspects clearly demonstrate the superiority of our sensing strategy over all gold standard DENV detection methods. Since dengue-infected patients can progress rapidly over a short period, early intervention may be life-saving. Current detection methods are able to detect infection only on or a few days after initial onset of symptoms. Our sensor, being able to directly and potently interact with the virus in serum and plasma, is further practically valuable as it would enable timely and earlier detection. Therefore, the sensor could serve as a rapid precautionary screening tool to control pandemic outbreaks, particularly in low-resource settings. Therapeutically, since negatively-charged host cell surface glycosaminoglycans (GAGs) widely interact with various pathogen surface proteins to facilitate host cell invasion in various infectious diseases (e.g., malaria parasite^36^, DENV^37^, ZIKV,^38^ and influenza A^14^), the negatively charged DNA nanoarchitecture developed herein likely confers additional viral inhibition effects in blood serum and plasma. Therefore, the overall inhibitory effect is achieved through specific, direct multivalent interactions with viral epitopes as well as general electrostatic interactions that allow DNA to strongly bind the virus and exhibit repulsion to the host cell surface GAGs, physically trapping and electrostatically isolating dengue virions from host cells.

## RESULTS AND DISCUSSION

### Design and characterization of star-shaped DNA architecture

Dengue virus (DENV, Type-2 strain), an enveloped arbovirus from the *Flaviviridae* family,^39^ was used herein to demonstrate our design strategy/concept. DENV was chosen because its envelope protein binding-domain III (ED3), a viral surface epitope, is organized into a complex icosahedral shape^40^ with alternating clusters of three or five ED3 sites (Fig. 1a); this represents an extremely challenging pattern to target using previous inhibitor design strategies. By connecting the clusters of ED3 sites linearly, we determined that a star-shape, consisting of a interior pentagon connected to five exterior triangles, would provide an optimal multivalent scaffold (Fig. s1b). This star design not only mirrors the global pattern of ED3 clusters (aligning with the star’s ten vertices) but also matches its local-surface, inter-cluster spacing distance of 15-nm between adjacent, alternating trivalent and pentavalent ED3 clusters. Based on this structural information, we designed a star-shaped DNA nanostructure (called “DNA star” herein) that was assembled from 21 DNA oligonucleotides of programmed sequences (Figs. 1b and s1c). The DNA star contains ten 15-nm long double-stranded (dsDNA) external edges in which each inner vertex is comprised of a 4-arm junction. Each inner edge carries a single-stranded DNA (ssDNA) region that can form a hairpin (stem loop) structure with a 6-bp (base-pair) long stem. When the hairpin is completely converted into the ssDNA form, its fully stretched length is also 15-nm. As further elaborated below, these hairpins serve dual functions, providing molecular beacon-like motifs for viral detection and offering local structural flexibility of ssDNA regions to ensure binding and inhibition of viral targets under various solution conditions and temperatures.

**Figure 1.**
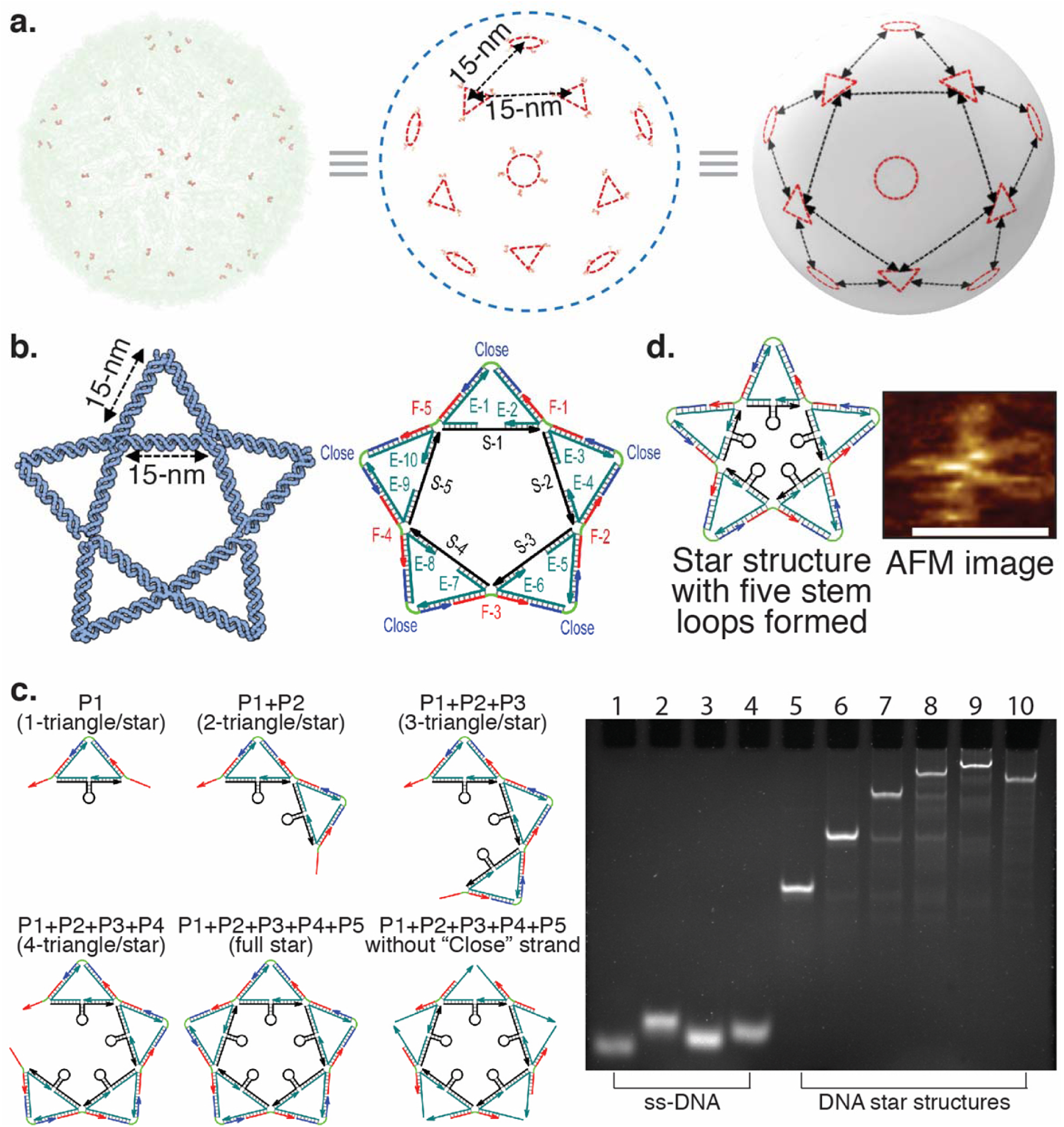
Design principle and characterization of a star-shaped DNA nanoarchitecture (DNA star). (**a**) Analysis of the pattern and spacing of DENV envelope protein domain III (ED3) clusters (shown as triangles or circles located ∼ 15 nm apart). (**b**) Based on the surface pattern of ED3 clusters, a star-shaped DNA nanostructure (DNA star) that contains five 4-arm junctions at the inner vertices is designed to match the pattern and spacing of ED3 clusters. Specifically, the DNA star is comprised of 21 oligonucleotides: five “scaffold” strands (S-1 to S-5) to form the internal edges, ten “edge” strands (E-1 to E-10) to connect internal and external edges, five “fix” strands (F-1 to F-5) to connect the external edges, and one “close” strand to cap the external edges of each triangle. (**c**) Characterization of the DNA star structure using native PAGE. On the left is the schematic of the DNA star structures characterized on the 4% non-denaturing PAGE. Lane-1: “Fix” strand; Lane-2: “Close” strand; Lane-3: “Scaffold” strand; Lane-4: “Edge” strand; Lane-5: P1; Lane-6: P1+P2; Lane-7: P1+P2+P3; Lane-8: P1+P2+P3+P4; Lane-9: P1+P2+P3+P4+P5 (full star); Lane-10: P1+P2+P3+P4+P5 (without “Close” strand). Based on the intensity quantifications of DNA species in each lane, yield of each full-size DNA complex formation (Lanes 6-10) is 100%, 98.2%, 81.2%, 86.3%, 88.1%, and 98.3%, respectively. (**d**) In the absence of viral target, single-stranded regions in S-1, S-2, S-3, S-4, and S-5 form stem loops. AFM image further confirms structural formation of DNA star in its “contracted” form. Scale bar indicates 50-nm.

The formation of the DNA star was first characterized by a non-denaturing polyacrylamide gel electrophoresis (PAGE) that showed the expected species that corresponded to the full DNA star complex (Lane-9 on the gel in Fig. 1c); other individual strands (single stranded DNA oligonucleotides) and partial star complexes are included for size reference in the non-denaturing gel. Based on the intensity of DNA species quantification on the gel, the yield of DNA star formation was ∼ 90%. This is quite high considering the difficulties in achieving perfect stoichiometry among all the component DNA strands. Atomic force microscopy (AFM) imaging further confirms formation of the designed DNA star nanostructure (Fig. 1d and Fig. s2). Note that the regular AFM probe may disturb individual DNA star structure during the scan. Additionally, AFM may display the overall shape of DNA star but cannot fully resolve the details of the structure due to the limited resolution.

### DNA star-based DENV sensing

A DNA star for viral detection and inhibition (below) of DENV (serotype 2, referred to as just DENV) was prepared by placing a ED3-binding aptamer (called “aptamer” herein)^41^ at all ten vertices of the DNA star to form a star-aptamer complex that geometrically matches and targets ED3 clusters (Fig. 2a). DENV was chosen since the aptamer is well characterized and available for this serotype. The aptamer is a weak binder to DENV surface with a *K*_*D*_ (dissociation constant) of ∼ 20 µM as determined by surface plasmon resonance (SPR) (Fig. s3). A FAM (fluorophore) or BHQ-1 (quencher) molecule-carrying ssDNA was hybridized to each inner edge strand flanking the hairpin to turn the star-aptamer complex into a viral sensor. This sensor design was based on our hypothesis that in the absence of DENV, FAM and BHQ-1 molecules would be brought together by Watson-Crick base pairing in the stem, much like a molecular beacon.^42^ Unlike a molecular beacon, which gives a fluorescent readout of target nucleic acid hybridization events, the hairpins would be pulled apart and converted to ssDNA (single-stranded DNA) as the aptamers arranged on the DNA star bind to ED3 sites on the DENV surface. The potent multivalent interactions, promoted by the matched geometric aptamer-ED3 pattern, would therefore cause a separation of the FAM fluorophores from BHQ-1 quenchers to afford a fluorescent readout. The ssDNA region also offers local structural flexibility, allowing for retention of equivalent binding ability under perturbations by matching the DNA star-templated aptamer pattern with slightly deformed arrangement of ED3 clusters due to temperature changes.^43^

**Figure 2.**
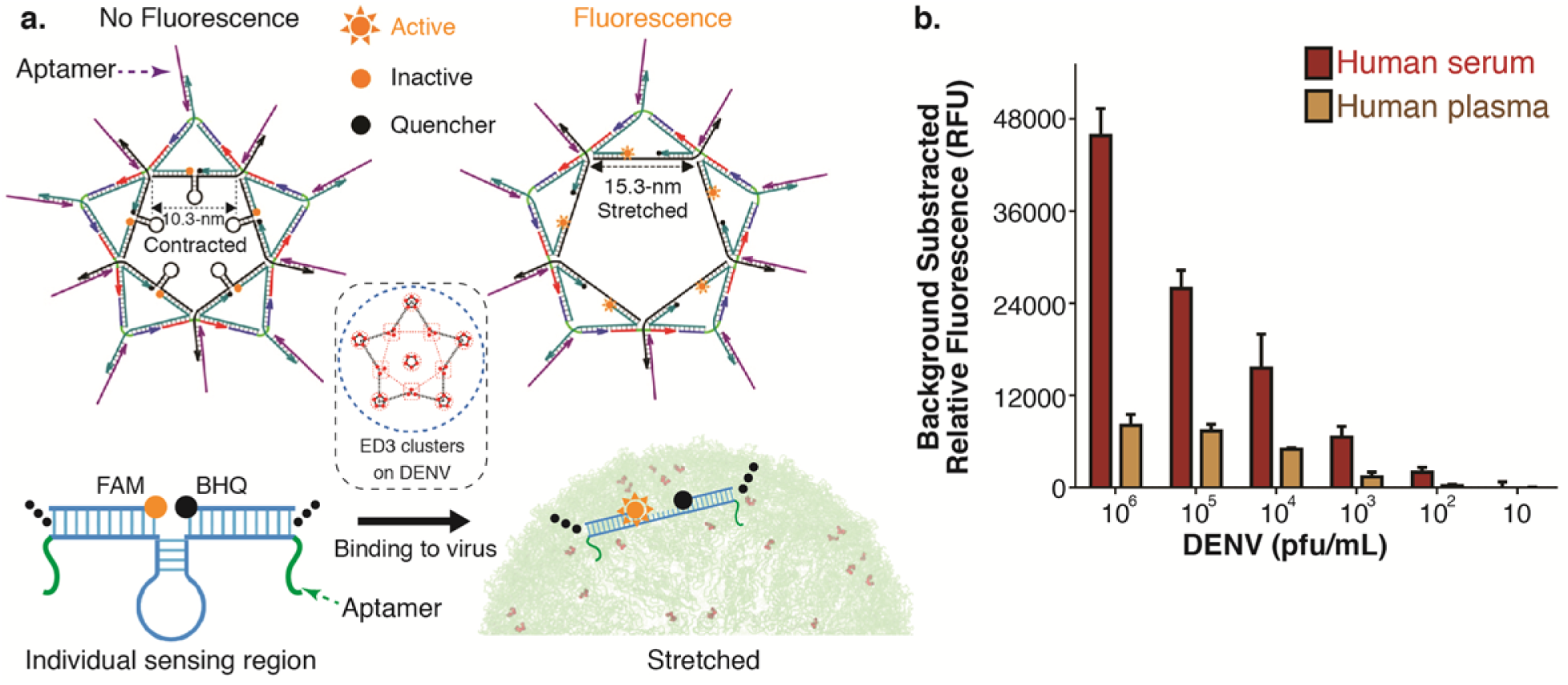
DNA star sensor. (**a**) The DNA star structure with 10 aptamers is made to match the pattern and spacing of envelope protein domain III (ED3) clusters on the DENV viral surface. Five molecular beacon-like sensing motifs, FAM-BHQ (fluorophore-quencher) pairs on all inner edges, are arranged in a quenching FRET by DNA base pairing in the hairpins. In the presence of DENV, the binding interactions between aptamers and ED3 domains would pull FAM away from BHQ by fully converting hairpins to ssDNA, enabling a fluorescent readout. Sensing motifs were designed to be flanked by trivalent ED3 clusters in the presence of DENV. (**b**) A series of DENV concentrations are used to determine the limit of detection (LOD) of the star sensor in human serum (100 pfu/mL) and human plasma (1,000 pfu/mL). The detection sensitivity in plasma is lower than that in serum due to greater amount of clotting proteins, leading to solution turbidity and blocking of fluorescence signal. Error bars are mean ± s.d., n = 3.

The experimental data support our hypothesis, showing that the DNA star-aptamer sensor was able to directly detect DENV virions with high sensitivity, affording a limit of detection (LOD) of 100 pfu/mL and 1,000 pfu/mL, respectively in human blood serum and plasma (Fig. 2b). Since plasma contains a greater amount of proteins, the solution exhibits some turbidity. This can prevent detection of FAM signal by obfuscation or scattering of light, leading to a lower sensitivity in plasma compared to serum. Additionally, our sensing platform is a simple mix-and-read assay that takes less than 5 min; the signal can be read by a portable fluorometer such as the Qubit Fluorometer from Life Technologies (excitation filters are 430-495 nm and 600-645 nm; emission filters are at 510-580 nm and 665-725 nm); and the cost for each sensing solution is only ∼ $0.15 (Note s2). The LOD of our sensor in serum or plasma is below or similar to the viral load level in patients between day [−2] and day [−1] (day [0] marks the onset of symptoms/illness which arises 3 days post infection); such high sensitivity cannot be achieved by the gold standard methods. The only two gold standard methods that can sense the infection of patients before the “onset of symptoms” (between day [−1] and day [0]) are viral isolation and RNA detection. However, both methods are time-consuming (1-2 weeks for viral isolation; 1-2 days for nucleic acid detection), expensive, and require a sophisticated clinical laboratory setup and technical expertise. ELISA-based antigen detection of NS1 is the next earliest and quickest to detection results of the gold standard methods, but it is expensive (hundreds of dollars) and cannot confirm the infection before the onset of symptoms as NS1 antigens are not available in high enough quantity. Additionally, the IgM ELISA or IgM rapid test cannot provide early viral sensing as IgM antibodies are only detectable in 50%, 80%, or 99% of patients 3-5 days, 5 days, or 10 days post infection. So, no gold standard method for DENV detection can achieve the same sensing capacity as that of our DNA star sensor in regard to the sensitivity, ease, quickness, and cost (see Fig. s4 and Table s2 for the detailed comparison). Additionally, our direct viral detection method can avoid the false positive sensing of IgG that is produced from the first DENV infection. In summary, the LOD of our sensor is well below the viral concentration (>10^5^ pfu/mL, see Note s2) in patients on day [0] (or the onset of illness, when fever and a variety of symptoms start to occur and the virus begins to become very pathogenic).^44^ Thus, detection before this date is highly beneficial to the patient and to the timely screening and control of pandemic outbreaks within surveillance and diagnostic networks. Given the ease, quickness, and low-cost of our sensing assay, our DNA star-based sensor would also be valuable in a low-resource or field-applicable environment.

### DNA 9-loop based sensing (a control sensor)

We next prepared a control DNA platform that precisely orchestrated the valency and spacing of a greater number of ED3-binding aptamer ligands and FAM-BHQ-1 pairs in 1D space to test our hypothesis that the previously determined highly sensitive DENV detection provided by our DNA star design must rely on 2D aptamer-ED3 pattern matching. As illustrated in Fig. 3a, a 6-helix bundle DNA origami nanotube (DNA-NT) carrying nine ssDNA loops (each 63-nt long on one side) and hinge points (on the other side) was designed using caDNAno (Fig. s5).^45^ Each ssDNA loop was flanked by a pair of FAM-BHQ molecules through DNA hybridization. Due to the stacking interactions (via π–π stacking) between adjacent blunt-ended DNA segments,^46–48^ we predicted that this bendable DNA-NT (called “9-loop” hereafter) would remain straight in the absence of binding targets and, thus, hold FAM and BHQ-1 molecules together in a quenching FRET. AFM imaging of the native 9-loop confirms this prediction by showing that the construct does remain straight with the expected length (Figs. 3a and s6). AFM results also show that multiple 9-loop structures can join together as a result of blunt-ended DNA stacking. If the binding affinity/force resulting from the binding of 9-loop-aptamer complex to spherical-shaped DENV was comparable to that of star-aptamer complex to DENV, the linear 9-loop should bend its segments about the hinge points to maximally bind ED3 sites (Fig. 3a). As a result, all the paired fluorophores and quenchers would become sufficiently separated to enable a fluorescence signal.

**Figure 3.**
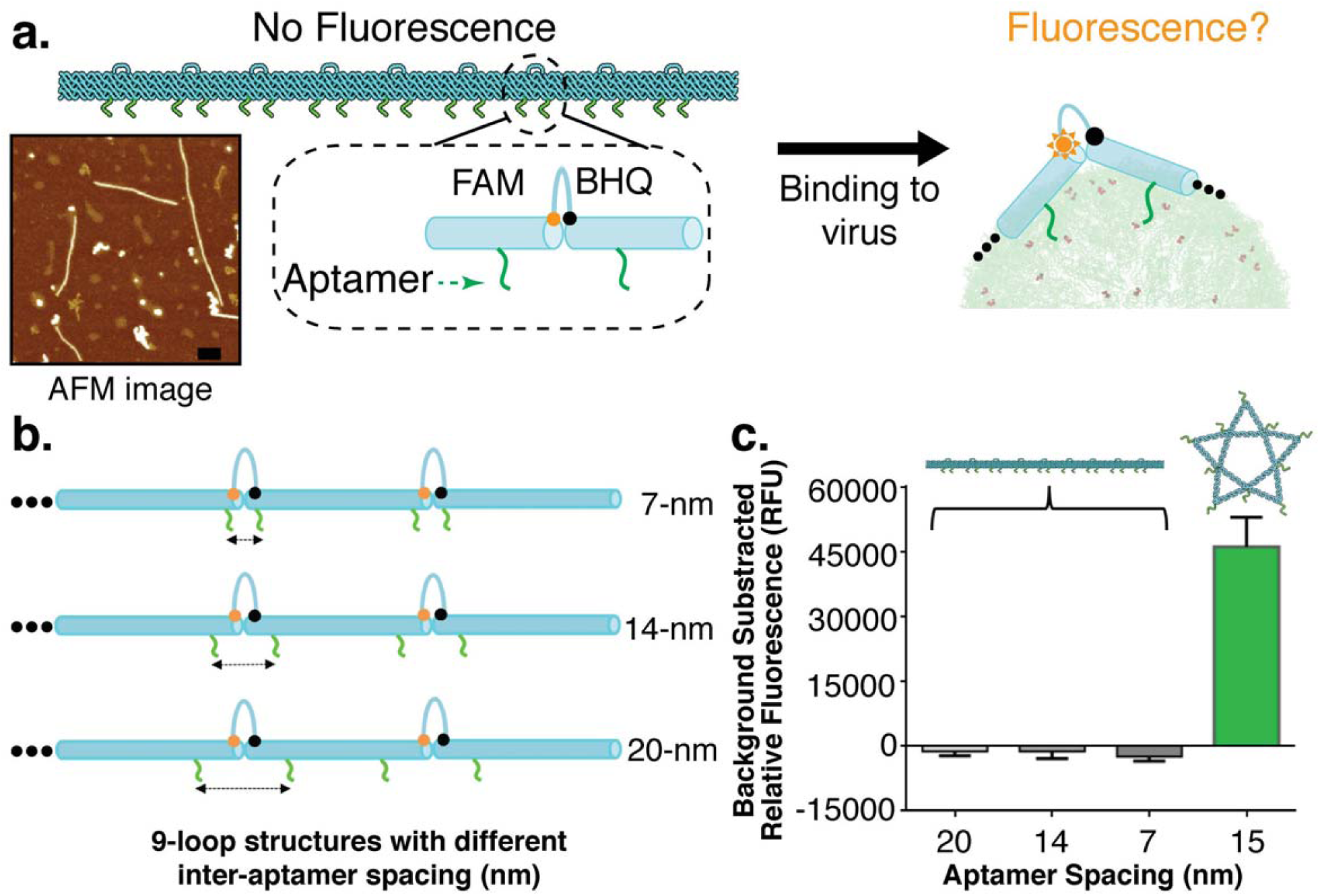
DNA 9-loop sensor. (**a**) The 9-loop templated DENV sensor contains 10 bendable segments linked by nine ssDNA loops (63-nt long on one side) and hinge points (on the other side). The light blue cylinders represent tubular-shaped 6-helix DNA bundles. AFM imaging of the 9-loop proves the structure remains straight at the correct length. The scale bar indicates 200 nm. FRET between FAM and BHQ occurs through blunt-ended DNA base stacking interactions between adjacent cylinders. Aptamer ligands are arranged on the 9-loop with different inter-ligand spacing (7, 14, or 21 nm). FAM quenching by BHQ would disappear if 9-loop-aptamer complex could interact with DENV with high avidity to pull FAM and BHQ apart. (**b**) Schematic of the 9-loop-aptamer complexes with 7-nm, 14-nm, and 20-nm inter-aptamer spacing. (**c**) The sensing capacity of the 9-loop or DNA star sensor is measured based on relative fluorescence units (RFU), demonstrating that the star sensor is able to detect DENV while the fluorescence of all 9-loop sensors is still quenched in the presence of DENV with the viral load of 10^7^ pfu/mL (See Note S6 for possible reason why the 9-loop sensing signal is negative). Error bars are mean ± s.d., n = 3.

We then decorated the 9-loop with eighteen ED3-binding aptamer ligands with various ligand spacing (7, 14, or 21 nm) in 1D space to test the ability of the 9-loop sensor to detect DENV (Fig. 3b). Our experimental data revealed that none of these sensors were able to detect the presence of DENV (Fig. 3c) even at a very high viral load of 10^7^ pfu/mL. Breaking the more stable Watson-Crick base pairing in the DNA star’s five hairpins is much harder than disturbing blunt-ended DNA stacking in the 9-loop construct. Therefore, our results suggest that the 2D spatial pattern-recognition display was an essential “element” for turning a weak-binding ligand into a significantly stronger binder that could be used for both detection and viral inhibition by physically binding and wrapping the viral particles (below). For DENV, simply matching the spacing in 1D is insufficient for converting weakly-binding ligands into a strong multivalent binder.

We further verified the 9-loop nanostructure’s sensing capabilities by using very strong biotin-streptavidin interactions to eliminate the possibility that the poor performance of the 9-loop-aptamer sensor resulted from an unexpected design flaw. For this purpose, 9-loop-biotin complexes were made to test whether the structure could offer designed sensing capacity by targeting streptavidin-coated magnetic beads. The 9-loop-biotin complexes mimic the 9-loop-aptamer sensors but relies on the much stronger (pM *K*_*D*_) interaction of biotin to streptavidin to validate the design.^49^ Since streptavidin spacing on magnetic beads is not known, various inter-biotin distances were tested to inclusively survey the sensing capacity of the 9-loop sensor (Fig. s7a). If biotin spacing matched the streptavidin spacing, the strong biotin-streptavidin interactions would bend the 9-loop segments flanking the ssDNA loops, moving the quenchers and fluorophores apart, abrogating quenching and resulting in direct fluorescence detection (Fig. s7b). Our data show that the fluorescence difference between quenched and unquenched states at different temperatures was significant; a biotin spacing of 7-nm flanking the loops afforded the best detection sensitivity (Figs. S8a/b). When biotin was spaced at a high density, at every 7-nm along the 9-loop, sensing capability was hindered due to steric hindrance from biotin crowding. In other words, a higher ligand density does not improve sensing capability even at the optimal inter-ligand spacing. This observation was further confirmed by fluorescence confocal microscopy image analysis (Fig. s8c). We observed that the detection sensitivity increased as the temperature decreased, consistent with blunt-ended DNA stacking being more stable at lower temperatures, decreasing spontaneous movement of fluorophores away from quenchers. Additionally, we observed that fluorescence intensity generally decreased as the temperature increased, which agrees with previous studies.^50,51^ It should be noted that the above biotin-streptavidin interaction-mediated sensing result is not contradictory to our design concept but rather supports the notion that a precise, multivalent spatial pattern-recognition strategy should be able to turn any ligand, weak or strong, into an even stronger binder.

### Binding affinity assay of DENV to DNA-star, 9-loop, and monovalent aptamer

The binding avidities of the star-aptamer and the various 9-loop-aptamer complexes to DENV were assayed by SPR to determine whether the differentiated DENV detection ability of the DNA star and 9-loop sensors is due to their different binding avidities to the virus (Fig. s9). Monovalent aptamer was used as a baseline control. Partial star-aptamer complexes (containing 1, 2, 3, or 4 triangles of the full star) serve as another set of controls to demonstrate the necessity of the full DNA star-aptamer complex to achieve high viral-binding affinity. The SPR results show that the binding of the DNA star-aptamer complex to DENV was stronger than all the partial star-aptamer complexes (Fig. 4a); its binding was also > 100-fold stronger than the 9-loop-aptamer complexes or monovalent aptamer when normalized to the same concentration of aptamers (Fig. 4b). Such observations confirm that matching both shape/pattern in 2D space and inter-ED3 spacing is critical for achieving high binding avidity (Figs. 4c and s10).

**Figure 4.**
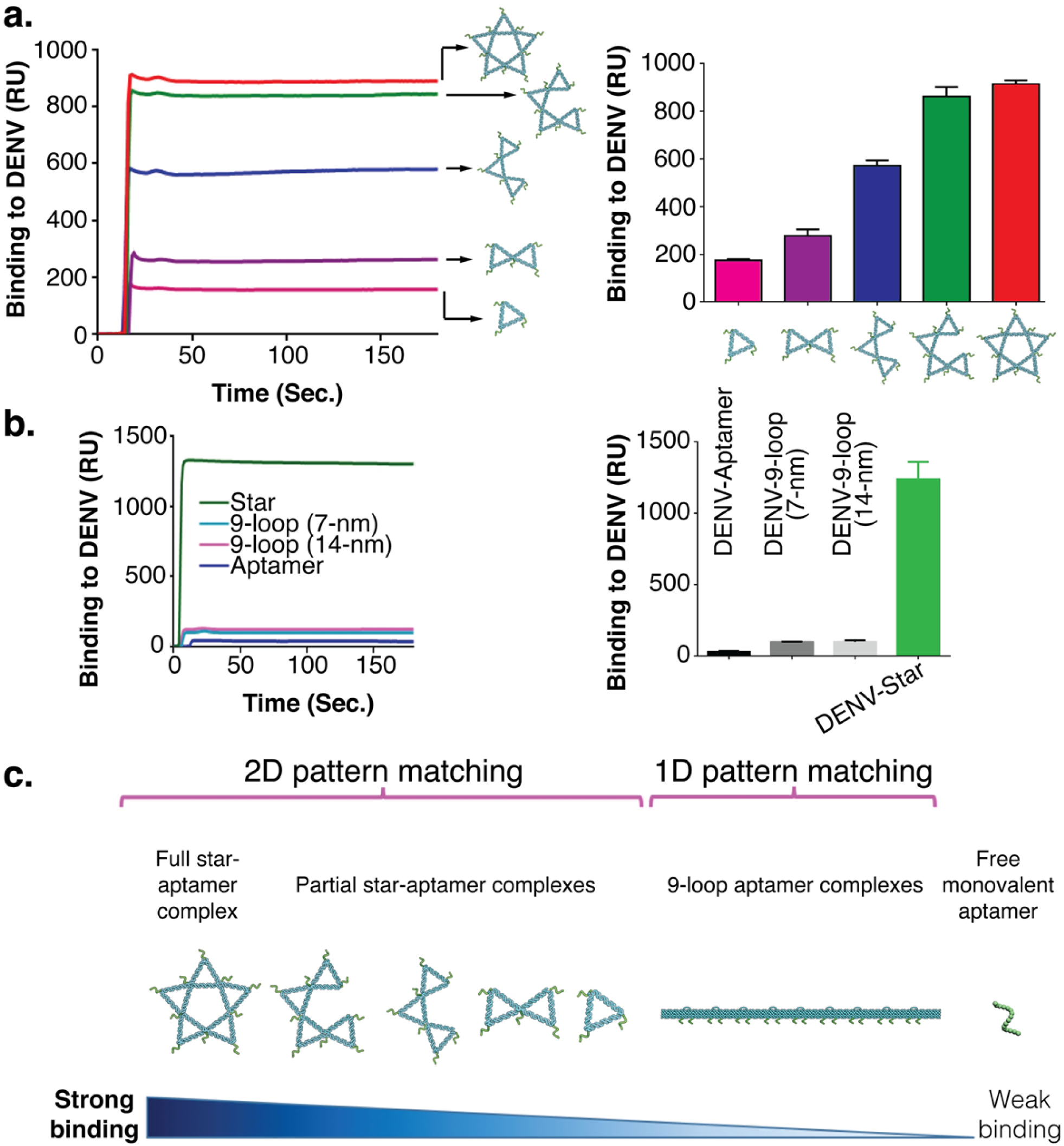
Multivalent binding assay using SPR. (**a**) Surface plasmon resonance (SPR) binding assays of the full star-aptamer complex and partial star-aptamer complexes with DENV at a normalized aptamer concentration of 1.20 µM. The full star-aptamer complex shows the strongest binding compared to all the partial star-aptamer complexes based on the relative SPR signal units. Error bars are mean ± s.d., n = 3. (**b**) SPR binding assays of free aptamer, two 9-loop-aptamer complexes, and star-aptamer complex with DENV at a normalized aptamer concentration of 1.8 µM. The star-aptamer complex shows ∼ 100-fold stronger viral binding compared to the 9-loop-aptamer complexes or free aptamer, based on the relative SPR signal units. Error bars are mean ± s.d., n = 3. (**c**) Side-by-side comparison of different DNA-based nanostructures’ binding strengths to DENV.

### *In vitro* DENV inhibition

Encouraged by the sensing and SPR results, we tested the *in vitro* inhibition of DENV (plaque forming and EC_50_ assays) using the DNA star-aptamer complex or monovalent aptamer (as a control) in human serum-containing solution optimized for a standard *in vitro* antiviral plaque assay. Briefly, the DENV serotype 2 viral particles were incubated with different concentrations of inhibitor (DNA star-aptamer complex or monovalent aptamer) in human serum and the remaining infectivity was determined by a plaque reduction assay (Fig. 5a). Next, the dose-dependent inhibition of DENV was examined. The EC_50_ value (half maximal effective concentration) of the DNA star-aptamer complex for DENV infection inhibition was ∼ 2 nM, whereas the EC_50_ value of the monovalent aptamer was ∼ 15 µM (Fig. 5b). These results demonstrate that the DNA star-aptamer multivalent inhibitor was ∼ 7.5 × 10^3^-fold more effective than the monovalent aptamer for the *in vitro* inhibition of DENV infection in human serum.

**Figure 5.**
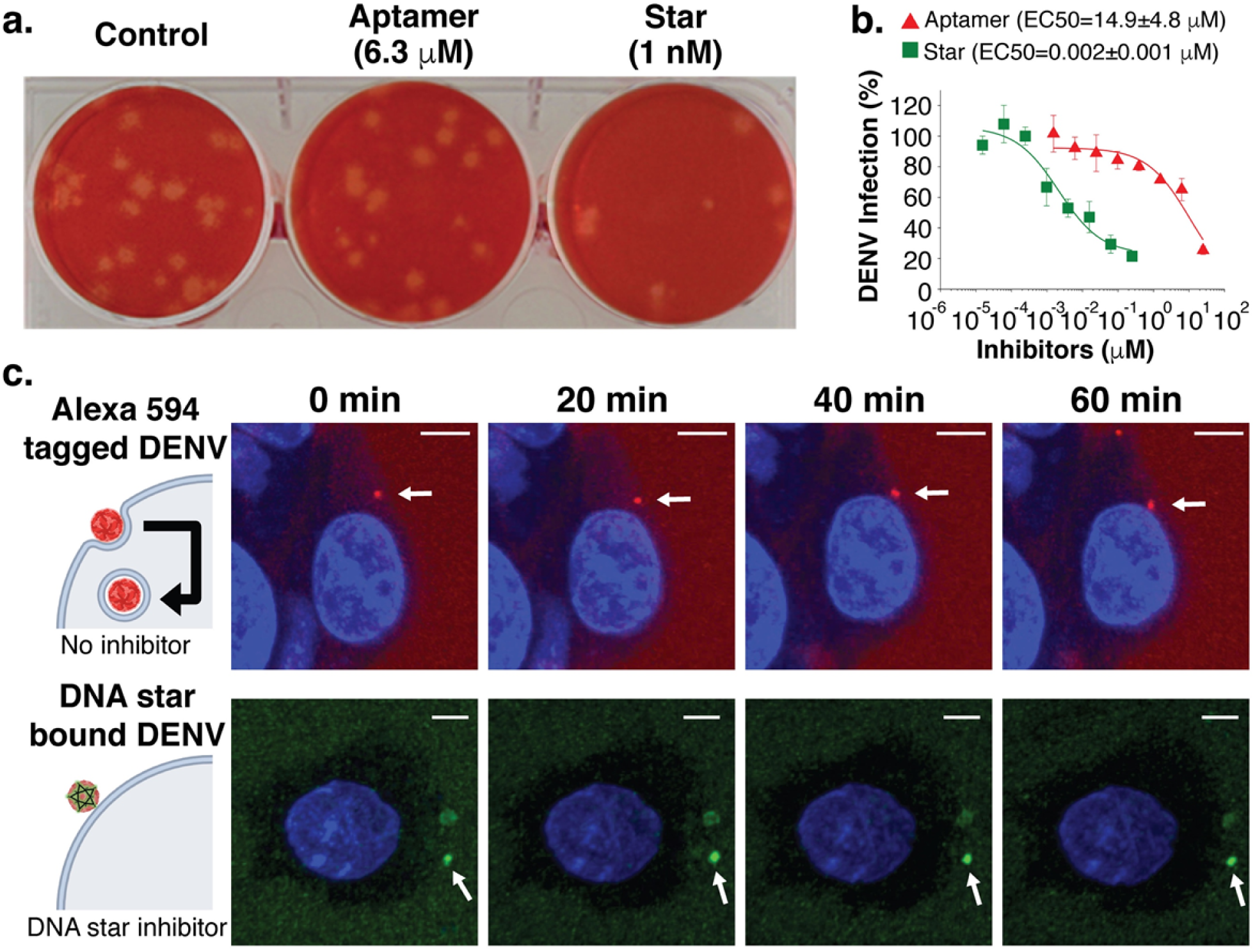
DENV inhibition profiling. (**a**) The plaque assay shows that the star-aptamer complex at a lower concentration of the aptamer (1 nM) can inhibit plaque formation better than the monovalent aptamer even at a higher concentration of the aptamer (6.3 µM). (**b**) EC_50_ assays show that the DNA star-aptamer complex is ∼ 7.5 × 10^3^-fold more effective than the monovalent aptamer at DENV infection inhibition in human blood serum. Error bars are mean ± s.e.m., n = 3. (**c**) Time-lapsed imaging assay of A594-labeled (**top row**) or DNA star sensor-bound DENV (**bottom row**) shows viral entry or inhibition on HepG2 cells. In the case of A594-labeled viruses (red dots), both cell binding and internalization are shown over time. DNA star sensor-bound viruses (green dots) do not show the cell internalization. Cell nuclei (blue color) were stained with Hoechst. For each case, a white arrow is used to track the virion that was in the plane of focus throughout the confocal imaging assay from time 0-min. The same virion for each case was tracked in the supplemental movies (Movies s1 and s2). Scale bars indicate 5-μm. Note that other virions (shown in Fig. s11 and Movies s1 and s2) were also imaged when they came in or out of the focal plane during the time course.

We then carried out a time-lapsed, live confocal imaging assay to demonstrate that DENV virions can lose their cell internalization/infection ability after being bound by the DNA star-aptamer complex. For the assay, HepG2 cells were mixed with the DNA star sensor-bound or Alexa 594-labeled virions in human serum-containing solution optimized for HepG2 cell culturing. HepG2 was chosen since it is a human cell line derived from the liver, an organ DENV can infect. The Alexa 594-labeled virion was used as a control to show positive viral cell entry/internalization. For each mixture, real-time imaging of the interactions between virions and HepG2 cells was performed. As shown in Fig. 5c, Fig. s11 and by Movie s1, after binding to DENV, the DNA star-aptamer complex is able to inhibit viral cell entry, and some of the DNA star-bound virions (green dots) also moved away from the HepG2 cells as time went on. On the contrary, the Alexa 594-labeled control virions (red dots) show the ability to adhere to the cell surface, and then pass through the cell membrane and approach the cell nucleus after internalization (Fig. 5c, Fig. s11 and Movie s2). In both conditions, HepG2 cell nuclei (blue color) were stained with Hoechst. Confocal imaging has confirmed that our DNA star-aptamer complex has the capability to inhibit DENV at a single viral particle level by physical trapping and isolating DENV from host cells.

## CONCLUSION AND OUTLOOK

Through targeting one of the most challenging exemplar platforms, the DENV flavivirus, we have demonstrated the unique concept of integrating a structurally defined DNA nanoarchitecture with precise, multivalent spatial pattern-recognition properties for direct, highly-sensitive, easy, and rapid viral detection. Our sensor has clearly shown superiority over gold standard DENV detection methods and has promising use in field-applicable or low-resource environments. As is standard with viral infection screening, secondary confirmation of infection by some of the standard methods (e.g., viral isolation or nucleic acid sensing) should still be employed after the initial precautionary screening in order to make a full documentation and evaluation of an epidemic outbreak. Moreover, other monovalent aptamer combinations have been used as antiviral agents for human viruses.^52^ Thus, our synthetic DNA star-aptamer complex has the potential to serve as a highly potent anti-DENV inhibitor in the treatment of infected individuals.

As binders specific to other pathogenic domains can be evolved or synthesized, our strategy could be widely applicable by following a general DNA nanoplatform design principle (illustrated in Fig. 6) as various DNA 2D/3D structures can be designed and synthesized on-demand to mimic simpler surface epitope patterns of pathogenic viruses, bacteria, and microbial toxins. Although DNA nanostructures have shown certain stability *in vivo*,^27,35^ the DNA star’s component strands may be chemically modified (e.g., through methylation) to further improve its stability when preparing *in vivo* antiviral assays. In summary, the reported design strategy, by providing a unique perspective on the precise spatial pattern-recognition of pathogenic epitopes, holds promise for broad and effective applications.

**Figure 6.**
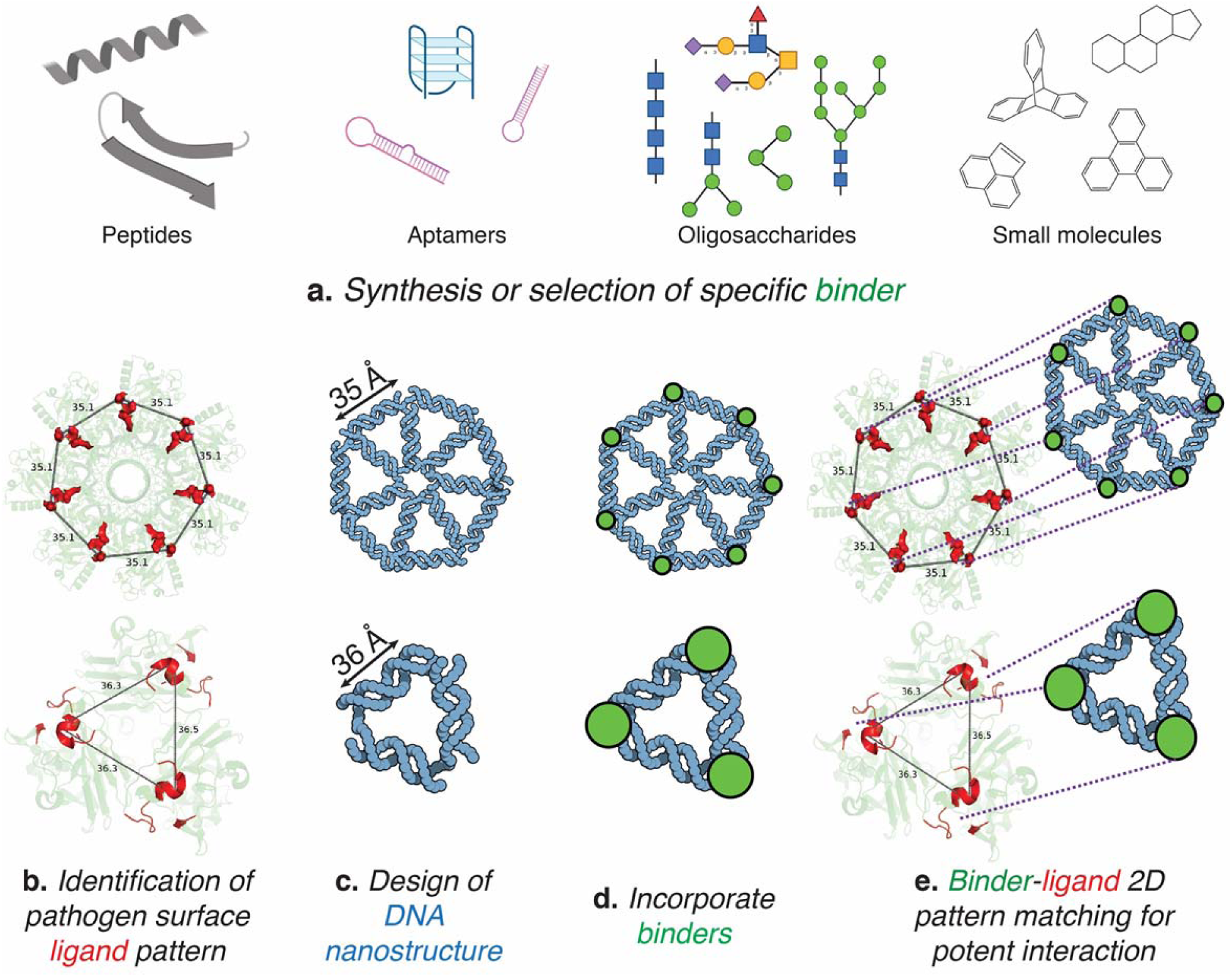
Design principle of DNA nanostructure-based pathogenic sensors and inhibitors. (**a**) Synthesize or evolve a specific binder against a pathogenic domain. Binders can include but are not limited to peptides, aptamers, oligosaccharides, or small molecules. (**b**) Analyze the arrangement to identify the spatial surface pattern of the pathogenic domain. (**c**) Design a 2D DNA nanostructure that mirrors the spatial surface pattern. (**d**) Incorporate binders at appropriate locations onto the 2D DNA nanostructure to match the pathogenic domain placement and spacing. (**e**) Adapt the potent, multivalent DNA nanostructure-binder complex to bind the pathogen for sensing or inhibition. Two simpler examples are used herein for illustration. **Top**: A 7-arm junction centered, heptagon-shaped DNA structure matches heptavalent binding sites on anthrax. **Bottom**: A triangle-shaped DNA structure matches hemagglutinin (HA) trimers found on the surface of influenza virus.

## ACKNOWLEDGMENTS

We thank the core facilities at RPI-CBIS, Wadsworth Center of New York State Department of Health, and Organic Electronics and Information Displays and Jiangsu Key Laboratory and Institute of Advanced Materials of Nanjing University of Posts and Telecommunications (China). Ahmad Abu-Hakmeh and Leo Wan are acknowledged for technical assistance with operating the StepOnePlus™ Real-Time PCR System.

## Author contributions

S-J.K., J.S.D., R.J.L. and X.W. conceived the original idea of employing designer DNA architecture for viral sensing and inhibition. For this specific article, P.S.K., S.-J.K. and X.W. designed and planned the experiments; S.R. and J.C. designed the sequences of DNA star, and performed non-denaturing PAGE and AFM experiments of DNA star; M.K. designed DNA 9-loop structure, prepared all the DNAs for AFM, EC_50_, plaque forming and time-lapsed confocal assays; L.K. and L.D.L. prepared viruses, and performed EC_50_ and plaque forming assay; F.Z. and N.C.S. contributed to the AFM experiments of DNA 9-loop structure; F.Z. performed SPR analysis; D.K. performed confocal microscopy analysis; K.F. analyzed the aptamer binding pattern of DENV; all authors contributed to the data analysis and manuscript preparation.

## Supplementary Information

Materials, Methods, Figs. s1 to s11, Tables s1 to s5, Notes s1 to s7, Movies s1 and s2, and References.

## References

1. Storch GA. Diagnostic virology. Clin Infect Dis 2000, 31(3): 739–751.

2. Read SJ, Burnett D, Fink CG. Molecular techniques for clinical diagnostic virology. J Clin Pathol 2000, 53(7): 502–506.

3. Scof S. Recent advances in diagnostic testing for viral infections. Biosci Horizons 2016, 9(1): 1–11.

4. Dengue: Guidelines for diagnosis, treatment, prevention and control: New edition: Geneva, 2009.

5. Whitehead SS, Blaney JE, Durbin AP, Murphy BR. Prospects for a dengue virus vaccine. Nat Rev Microbiol 2007, 5: 518–528.

6. Boni MF. Vaccination and antigenic drift in influenza. Vaccine 2008, 26(18): C8–C14.

7. Harrison SC. Virology. Looking inside adenovirus. Science 2010, 329(5995): 1026–1027.

8. Krantz BA, Melnyk RA, Zhang S, Juris SJ, Lacy DB, Wu Z, et al. A phenylalanine clamp catalyzes protein translocation through the anthrax toxin pore. Science 2005, 309(5735): 777–781.

9. Bandlow V, Liese S, Lauster D, Ludwig K, Netz RR, Herrmann A, et al. Spatial screening of hemagglutinin on influenza a virus particles: Sialyl-lacnac displays on DNA and peg scaffolds reveal the requirements for bivalency enhanced interactions with weak monovalent binders. J Am Chem Soc 2017, 139(45): 16389–16397.

10. Mourez M, Kane RS, Mogridge J, Metallo S, Deschatelets P, Sellman BR, et al. Designing a polyvalent inhibitor of anthrax toxin. Nat Biotechnol 2001, 19(10): 958–961.

11. Rai P, Padala C, Poon V, Saraph A, Basha S, Kate S, et al. Statistical pattern matching facilitates the design of polyvalent inhibitors of anthrax and cholera toxins. Nat Biotechnol 2006, 24(5): 582–586.

12. Joshi A, Punyani S, Bale SS, Yang H, Borca-Tasciuc T, Kane RS. Nanotube-assisted protein deactivation. Nat Nanotechnol 2008, 3(1): 41–45.

13. Kitov PI, Sadowska JM, Mulvey G, Armstrong GD, Ling H, Pannu NS, et al. Shiga-like toxins are neutralized by tailored multivalent carbohydrate ligands. Nature 2000, 403(6770): 669–672.

14. Kwon SJ, Na DH, Kwak JH, Douaisi M, Zhang F, Park EJ, et al. Nanostructured glycan architecture is important in the inhibition of influenza a virus infection. Nat Nanotechnol 2017, 12(1): 48–54.

15. Ahmad KM, Xiao Y, Soh HT. Selection is more intelligent than design: Improving the affinity of a bivalent ligand through directed evolution. Nucleic Acids Res 2012, 40(22): 11777–11783.

16. King DJ, Noss RR. Toxicity of polyacrylamide and acrylamide monomer. Rev Environ Health 1989, 8(1-4): 3–16.

17. Malik N, Wiwattanapatapee R, Klopsch R, Lorenz K, Frey H, Weener JW, et al. Dendrimers: Relationship between structure and biocompatibility in vitro, and preliminary studies on the biodistribution of 125i-labelled polyamidoamine dendrimers in vivo. J Control Release 2000, 65(1-2): 133–148.

18. Strauch EM, Bernard SM, La D, Bohn AJ, Lee PS, Anderson CE, et al. Computational design of trimeric influenza-neutralizing proteins targeting the hemagglutinin receptor binding site. Nat Biotechnol 2017, 35(7): 667–671.

19. Lin C, Liu Y, Rinker S, Yan H. DNA tile based self-assembly: Building complex nanoarchitectures. Chemphyschem 2006, 7(8): 1641–1647.

20. Chandrasekaran AR, Anderson N, Kizer M, Halvorsen K, Wang X. Beyond the fold: Emerging biological applications of DNA origami. Chembiochem 2016, 17(12): 1081– 1089.

21. Hong F, Zhang F, Liu Y, Yan H. DNA origami: Scaffolds for creating higher order structures. Chem Rev 2017.

22. Hu Q, Li H, Wang L, Gu H, Fan C. DNA nanotechnology-enabled drug delivery systems. Chem Rev 2018.

23. Seeman NC. Nucleic acid junctions and lattices. J Theor Biol 1982, 99(2): 237–247.

24. Zhang Q, Jiang Q, Li N, Dai LR, Liu Q, Song LL, et al. DNA origami as an in vivo drug delivery vehicle for cancer therapy. ACS Nano 2014, 8(7): 6633–6643.

25. Lee DS, Qian H, Tay CY, Leong DT. Cellular processing and destinies of artificial DNA nanostructures. Chem Soc Rev 2016, 45(15): 4199–4225.

26. Vinther M, Kjems J. Interfacing DNA nanodevices with biology: Challenges, solutions and perspectives. New J Phys 2016, 18: 085005.

27. Li S, Jiang Q, Liu S, Zhang Y, Tian Y, Song C, et al. A DNA nanorobot functions as a cancer therapeutic in response to a molecular trigger in vivo. Nat Biotechnol 2018, 36(3): 258–264.

28. Mei QA, Wei XX, Su FY, Liu Y, Youngbull C, Johnson R, et al. Stability of DNA origami nanoarrays in cell lysate. Nano Lett 2011, 11(4): 1477–1482.

29. Hahn J, Wickham SFJ, Shih WM, Perrault SD. Addressing the instability of DNA nanostructures in tissue culture. ACS Nano 2014, 8(9): 8765–8775.

30. Agarwal NP, Matthies M, Gur FN, Osada K, Schmidt TL. Block copolymer micellization as a protection strategy for DNA origami. Angew Chem Int Ed Engl 2017, 56(20): 5460– 5464.

31. Perrault SD, Shih WM. Virus-inspired membrane encapsulation of DNA nanostructures to achieve in vivo stability. ACS Nano 2014, 8(5): 5132–5140.

32. Rothemund PW. Folding DNA to create nanoscale shapes and patterns. Nature 2006, 440(7082): 297–302.

33. Wilner OI, Willner I. Functionalized DNA nanostructures. Chem Rev 2012, 112(4): 2528–2556.

34. Lee H, Lytton-Jean AK, Chen Y, Love KT, Park AI, Karagiannis ED, et al. Molecularly self-assembled nucleic acid nanoparticles for targeted in vivo sirna delivery. Nat Nanotechnol 2012, 7(6): 389–393.

35. Jiang D, Ge Z, Im HJ, England CG, Ni D, Hou J, et al. DNA origami nanostructures can exhibit preferential renal uptake and alleviate acute kidney injury. Nat Biomed Eng 2018, 2(11): 865–877.

36. Beaudet JM, Mansur L, Joo EJ, Kamhi E, Yang B, Clausen TM, et al. Characterization of human placental glycosaminoglycans and regional binding to var2csa in malaria infected erythrocytes. Glycoconj J 2014, 31(2): 109–116.

37. Chen Y, Maguire T, Hileman RE, Fromm JR, Esko JD, Linhardt RJ, et al. Dengue virus infectivity depends on envelope protein binding to target cell heparan sulfate. Nat Med 1997, 3(8): 866–871.

38. Kim SY, Zhao J, Liu X, Fraser K, Lin L, Zhang X, et al. Interaction of zika virus envelope protein with glycosaminoglycans. Biochemistry 2017, 56(8): 1151–1162.

39. Simmonds P, Becher P, Bukh J, Gould EA, Meyers G, Monath T, et al. Ictv virus taxonomy profile: Flaviviridae. J Gen Virol 2017, 98(1): 2–3.

40. Zhang X, Ge P, Yu X, Brannan JM, Bi G, Zhang Q, et al. Cryo-em structure of the mature dengue virus at 3.5-a resolution. Nat Struct Mol Biol 2013, 20(1): 105–110.

41. Chen HL, Hsiao WH, Lee HC, Wu SC, Cheng JW. Selection and characterization of DNA aptamers targeting all four serotypes of dengue viruses. PLoS One 2015, 10(6): e0131240.

42. Tyagi S, Kramer FR. Molecular beacons: Probes that fluoresce upon hybridization. Nat Biotechnol 1996, 14(3): 303–308.

43. Fibriansah G, Ng TS, Kostyuchenko VA, Lee J, Lee S, Wang J, et al. Structural changes in dengue virus when exposed to a temperature of 37 degrees c. J Virol 2013, 87(13): 7585–7592.

44. Gubler DJ. Dengue and dengue hemorrhagic fever. Clin Microbiol Rev 1998, 11(July): 480–496.

45. Douglas SM, Marblestone AH, Teerapittayanon S, Vazquez A, Church GM, Shih WM. Rapid prototyping of 3d DNA-origami shapes with cadnano. Nucleic Acids Res 2009, 37(15): 5001–5006.

46. Wang R, Kuzuya A, Liu W, Seeman NC. Blunt-ended DNA stacking interactions in a 3-helix motif. Chem Commun 2010, 46(27): 4905–4907.

47. Maffeo C, Luan B, Aksimentiev A.End-to-end attraction of duplex DNA. Nucleic Acids Res 2012, 40(9): 3812–3821.

48. Yakovchuk P, Protozanova E, Frank-Kamenetskii MD. Base-stacking and base-pairing contributions into thermal stability of the DNA double helix. Nucleic Acids Res 2006, 34(2): 564–574.

49. Green NM. Avidin. Adv Protein Chem 1975, 29: 85–133.

50. Liu WT, Wu JH, Li ES, Selamat ES. Emission characteristics of fluorescent labels with respect to temperature changes and subsequent effects on DNA microchip studies. Appl Environ Microbiol 2005, 71(10): 6453–6457.

51. Anderson N, Dinolfo PH, Wang X. Synthesis and characterization of porphyrin–DNA constructs for the self-assembly of modular energy transfer arrays. J Mater Chem C 2018, 6(10): 2452–2459.

52. Gonzalez VM, Martin ME, Fernandez G, Garcia-Sacristan A. Use of aptamers as diagnostics tools and antiviral agents for human viruses. Pharmaceuticals 2016,9(4).

